# Multi-omics Analysis of Human Blood Cells Reveals Unique Features of Age-associated Type2 CD8 Memory T cells

**DOI:** 10.64898/2025.12.23.696247

**Authors:** Hiroyuki Matsui, Marlene Cervantes, Mir M. Khalid, Alan Tomusiak, Varun Dwaraka, Jorge Landgrave-Gomez, Prasanna Vadhana Ashok Kumaar, Qingwen Chen, Jesica Lasky-Su, Jake Stone, Ritesh Tiwari, Ryan Kwok, Shuntaro Ichikawa, Benjamin D. Ambrose, Rebeccah R. Riley, Genesis Vega Hormazabal, Ariel Floro, Andreea Cristina Alexandru, Ryan Smith, Birgit Schilling, Herbert G. Kasler, Eric Verdin

**Author notes:** These authors contributed equally to this work (Hiroyuki Matsui, Marlene Cervantes, Mir M. Khalid).

## Abstract

Aging impacts immune function, but the mechanisms driving age-related changes in immune cell subsets remain unclear. To explore age-dependent changes in immune cell populations, we analyzed human peripheral blood mononuclear cells (PBMCs) from a cohort of healthy donors aged 20–82 years using a 36-color spectral flow cytometry panel focused on T cells. We identified a unique population of memory CD8 T cells, which lack CXCR3 and produce a Th2-like cytokine response, accumulate with age. We discovered an age-dependent bias in naïve CD8 T cells toward Th2 cytokine production, accompanied by transcriptional and epigenetic changes supporting this phenotype. Moreover, health outcome association analysis linked the accumulation of these unique CXCR3- central memory CD8 T cells to asthma, chronic liver conditions, and type 2 diabetes. Together, our results support the model that an age-dependent drift in epigenetic regulation towards a Th2-like phenotype drives a pathogenic Th2-like immune population.

## Introduction

Aging reshapes immune function, diminishing the capacity to effectively combat infections, respond robustly to vaccines, and regulate sterile inflammation. These age-related changes are driven by well-recognized shifts in immune cell composition and function, notably marked by decreased generation of naïve T cells due to thymic involution, which impairs immune repertoire diversity and promotes accumulation of dysfunctional T-cell populations in peripheral tissues^1^. In parallel, there is reduced production of naïve T and B cells due to bone marrow decline, along with decreased T-cell receptor (TCR) diversity in the peripheral naïve repertoire. Concurrently, memory and age-associated B cells expand, while myeloid cells increase in relative abundance, disrupting the balance between adaptive and innate immunity ^2–4^. These changes collectively impair immune responses and increase susceptibility to chronic inflammatory conditions, infections, and cancer.

Broad changes in T cell composition during aging have been well established, with significant declines in naïve T cells and a concomitant increase in polarized central and effector memory T cell subsets^5^. These memory subsets, which are clonally restricted and epigenetically primed to initiate essential rapid recall responses, are generated via cycles of antigen receptor activation in the contexts of different specific cytokine signals. These signals tightly regulate the epigenetic imprinting of different effector programs appropriate to specific pathogen challenges^6^. The first discovered cytokine-driven effector program switch was the Th1/Th2 axis, initiated through signaling via IL-12 and IL-4, respectively, which in turn induce transcription factors T-bet and GATA3^7^. T-bet and GATA3 then promote expression of the downstream effector molecules associated with each program (e.g. IFN-γ for Th1, IL-13 for Th2), while repressing genes of the opposing program^8^. This mutually antagonistic property contributes to both the stability and mutual exclusivity of these effector programs, maintaining tight regulation against inappropriate T-cell responses. However, differentiated CD4+ T cells retain a degree of epigenetic plasticity and can interconvert between Th1 and Th2 states under certain conditions^9^. In addition to the Th1/Th2 axis, other effector axes (e.g., Th17/Treg) have been defined, creating a complex landscape of immune subtypes whose balance is further perturbed with aging^10^.

Importantly, T cells with aberrant effector responses have been implicated in the pathogenesis of chronic inflammatory diseases such as asthma and allergy in the case of Th2 cells, and type 1 diabetes in the case of Th1 cells^11,12^. The risk of acquiring several such conditions also increases with age, suggesting that the T cell repertoire might become more prone to adopting aberrant effector responses in older individuals. Recent studies have also reported increased CD4+Th2 cells during aging, proposing that aging itself may exert an influence on T cell effector programming^13^. Cells that have adopted or are prone to adopt such fates should, in many cases, be identifiable by specific combinations of surface markers. However, the complexity of the different possible combinations that could define such populations has likely hampered their identification.

In this study, we addressed this lack of resolution by employing spectral flow cytometry, using a 36-color panel focused primarily on T cells, to detect nuanced, previously unrecognized changes in T-cell subsets during aging by providing a high-resolution, combinatorial view of markers and functional states. In this manner, we identified different subsets of CD8+ T cells increasing with age. Including CXCR3+ and CXCR3- CD8 central memory T cells (CXCR3- CM and CXCR3+ CM respectively). Surprisingly, CXCR3- CM exhibited transcriptional and epigenetic signatures associated with canonical Th2 memory function, including priming at genomic loci linked to Th2 cytokine production. Our findings reveal that aging may drive the epigenetic reprogramming of these CXCR3- CM toward a pathogenic state, contributing to chronic inflammatory conditions such as type 2 diabetes, chronic liver disease, and asthma. By uncovering these novel mechanisms, we provide new insights into how aging shapes immune dysregulation and offer potential therapeutic targets to mitigate its effects.

## Results

### Previously reported age-dependent changes in major PBMC lineages were also observed in our donor cohort

To elucidate the changes in cellular subsets within peripheral blood mononuclear cells (PBMCs) during aging, we analyzed total PBMCs from 45 young and older donors. Our study cohort was well-balanced with respect to sex and cytomegalovirus (CMV) serostatus (Fig. 1A). Using a 36-color reagent panel focused on markers of T cell differentiation, activation, and exhaustion (Supplementary Table 1), we first identified eight major PBMC subsets using conventional gating based on canonical lineage markers (Supplementary Fig. 1). These subsets were visualized using a UMAP plot, showing their unique distribution patterns (Fig. 1B).

**Figure 1.**
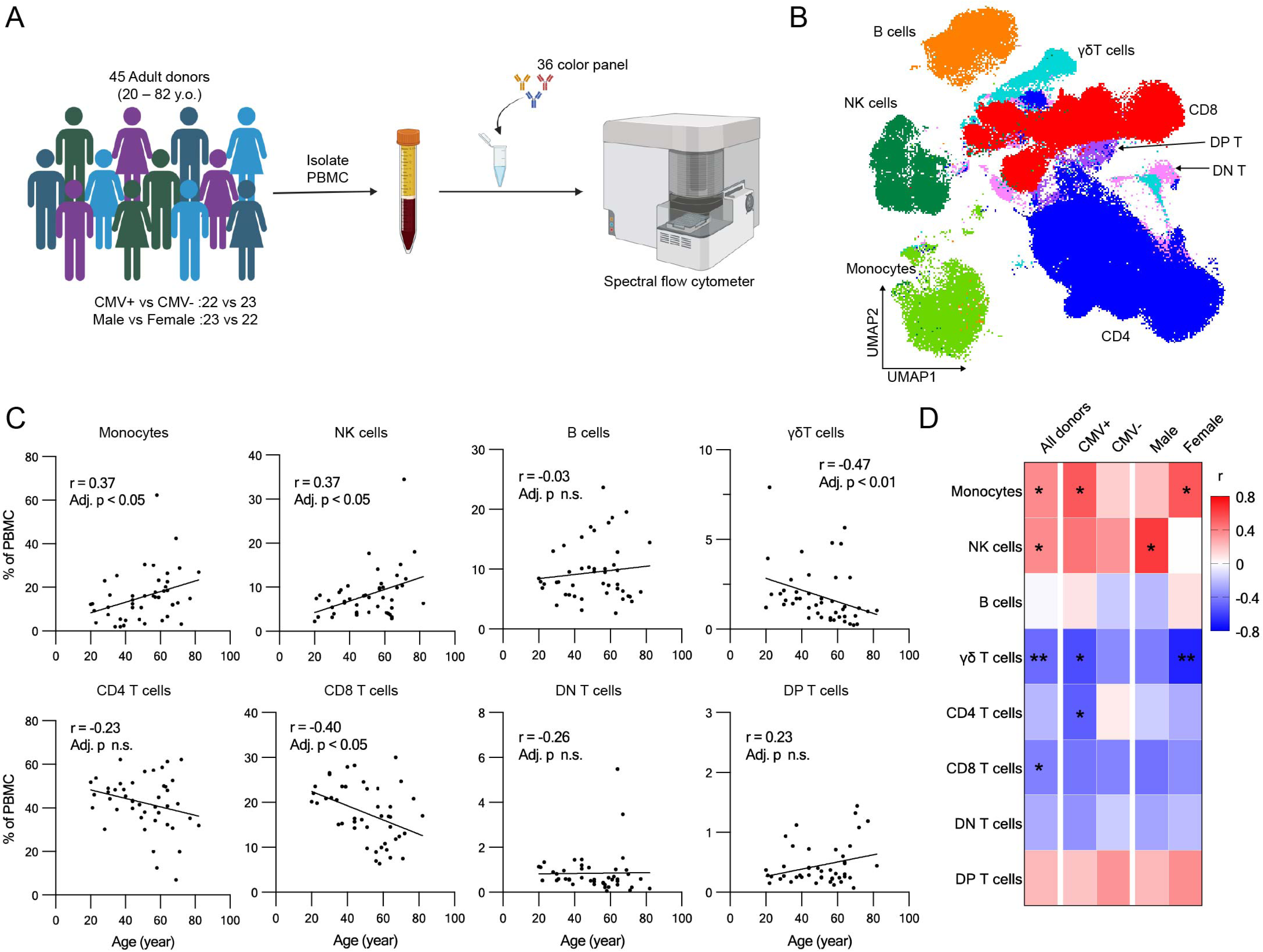
Cell composition changes during aging in human peripheral blood mono-nuclear cells. **A.** Schematic workflow of cell composition analysis with high dimensional spectral flow cytometer. **B.** UMAP plot of PBMCs from blood donors. **C.** Cell composition changes with age in individual PBMC subsets (n = 45, Spearman correlation test with Benjamini-Hochberg correction). **D.** Heatmap of correlation coefficient of age correlation analysis. Correlation coefficient of age correlation analysis in the individual subsets in the indicated donor groups are shown (n = 45, Spearman correlation test with Benjamini-Hochberg correction; * Adjusted p < 0.05, ** Adjusted p < 0.01).

Measuring the frequency of these subsets vs. donor age, we found a significant increase in monocytes and NK cells with advancing age, whereas CD8 T cells exhibited a significant decrease, as previously described^14–16^ (Fig. 1C). Additionally, γδ T cells showed a significant decline with age (Fig. 1C).

Analyzing subgroups based on sex and CMV serostatus, we found that monocytes increased more strongly with age in both CMV+ and female donors, while NK cells exhibited a stronger increase with age in male and CMV+ donors. Additionally, CD4 T cells significantly decreased in CMV+ donors, but were unchanged in the CMV- subgroup. In contrast, γδ T cells and CD8 T cells both showed similar, decreasing trends in all groups. These findings suggest that CMV serostatus and sex affect age-related changes in the frequency of monocytes, NK cells, and CD4 T cells, but not on CD8 or γδ T cells (Fig. 1D and Supplementary Fig. 2).

### Four CD8 T cell subsets showed significant changes during aging

Focusing on CD8 T cells, we performed FlowSOM clustering based on secondary markers and identified 12 subsets that occupy distinct clusters (Fig. 2A and B). These included the following: naïve CD8 T cells (CD45RA+CCR7+CD95-); two subsets of central memory CD8 T cells (TCM) (CD45RA-CCR7+) differentiated by the level of CXCR3 expression (CXCR+ and CXCR3-); three subsets of effector memory T cells (CD45RA-CCR7-) based on the expression of CD28, NKG2A and CD57 (CD28-CD57-NKG2A-, CD28+CD57-NKG2A+ and CD28+CD57+NKG2A-), four subsets of terminal effector memory CD8 T cells (CD45RA+CCR7-) distinct in their expression of CD57 and KIR (CD57-KIR-, CD57-KIR+, CD57+KIR- and CD57+KIR+), activated T cells with high expression of CD38, and mucosa-associated invariant T (MAIT), based on the expression of CD161 (Fig. 2B).

**Figure 2.**
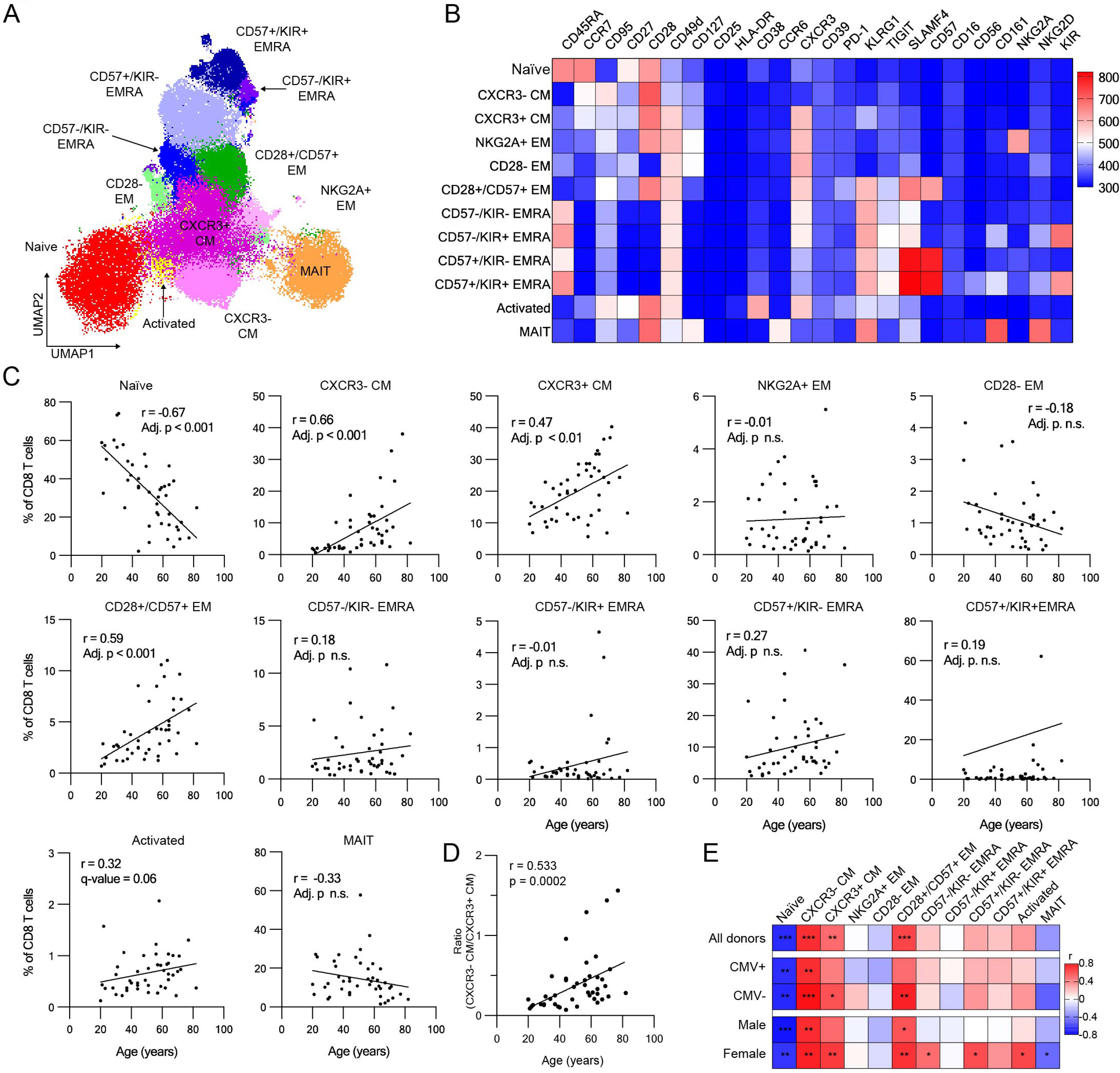
High-dimensional flow analysis with FlowSOM clustering revealed cell composition changes during aging in CD8 T cells. **A.** UMAP plot of CD8 T cells from blood donors. CD8 T cell subsets identified with FlowSOM clustering were shown in the different colors. **B.** Heatmap of the scaled fluorescent intensity of surface markers in individual subsets. **C.** Cell composition changes during aging in individual CD8 T cell subsets (n = 45, Spearman correlation test with Benjamini-Hochberg correction). **D.** Changes in the ratio of CXCR3+ CM over CXCR3- CM with age (n = 45, Spearman correlation test). **E.** Heatmap of correlation coefficient of age correlation analysis. Correlation coefficient of age correlation analysis in the individual CD8 T cell subsets in the indicated donor groups are shown (n = 45, Spearman correlation test with Benjamini-Hochberg correction; * Adjusted p < 0.05, ** Adjusted p < 0.01, *** Adjusted p < 0.001).

We then analyzed the correlation of the frequencies of these T cell subsets with age, both as a whole cohort and stratifying the data by sex and CMV status. As expected, naïve T cells showed a significant decrease with age (Fig. 2C), a trend consistently observed across all the donor subgroups (Fig. 2E, Supplementary Figs. 2 and 3). Both the CXCR3+ and CXCR3-CM subsets significantly increased with age (Fig. 3C), showing similar trends across all donor groups (Fig. 2E, Supplementary Figs. 2 and 3). We also identified CD28+/CD57+ EM as another subset showing a significant increase during aging (Fig. 2C). This subset exhibited significant changes in CMV- and in both male and female donors and a similar trend in CMV+ donors (Fig. 2E, Supplementary Figs. 2 and 3). Finally, CD57-/KIR- TEMRA, CD57+/KIR-TEMRA, activated T and MAIT cells exhibited significant increases exclusively in female donors (Fig. 2D, Supplementary Figs. 2 and 3). Of all these subsets, CXCR3- CM exhibited the most robust age-dependent increase across all donor groups, starting from a very low abundance in young donors and becoming increasingly abundant with respect to both total CD8 T cells and their CXCR3+ counterparts (Fig. 2D), highlighting their potential as key indicators of aging in CD8 T cells.

**Figure 3.**
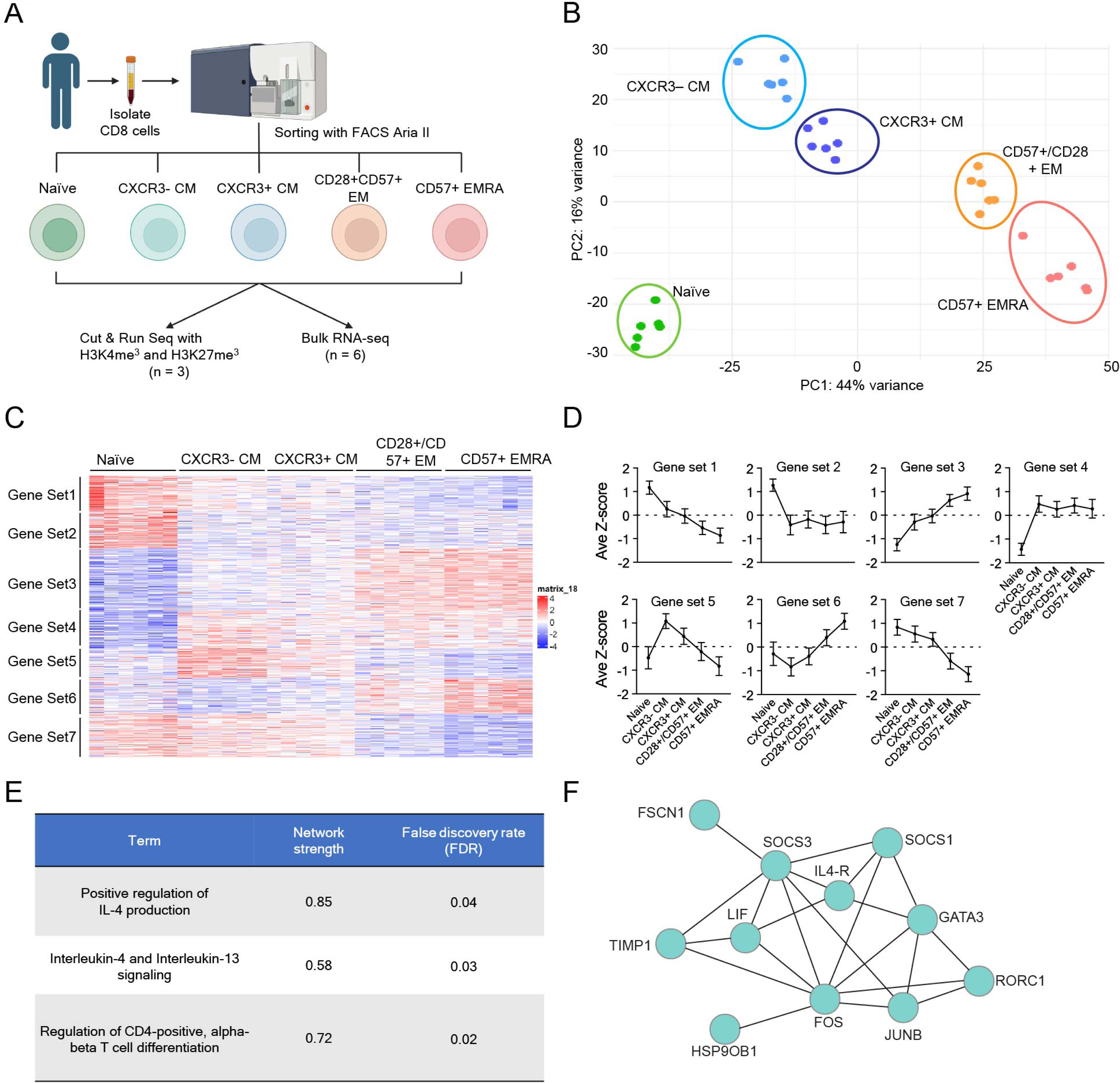
Transcriptomic analysis revealed CXCR3- CM has a distinct transcriptome profile and a feature of type-2 memory T cells. **A.** The schematic workflow of multi-omics analysis (RNA-seq, CUT & RUN with H3K4me3 and H3K27me3). Isolated CD8 T cells were sorted into 5 subsets (Naïve, CXCR3- CM, CXCR3+ CM, CD28+/CD57+ EM and CD57+ EMRA). After sorting, RNA extraction, CUT & were performed respectively. **B.** PCA plot of RNA-seq samples. Each dot shows individual samples. Each subset is shown in different colors. **C.** Heatmap of 6,471 differentially expressed genes across CD8 T cell subsets. Gene sets identified by clustering were shown on the left side. **D.** Average scaled expression levels of genes in individual gene sets across T cell subsets (average z-score ± SD). **E.** Significantly enriched terms in the Functional enrichment analysis by using gene set 5 using STRING database. **F.** Network of the genes included in the gene set 5 among Interlekin-4 and Interluekin-13 signaling pathway.

### CXCR3- CM shows a transcriptional and epigenetic profile resembling Th2 cells

We identified four age-dependent CD8 T cell subsets: naïve, CXCR3- CM, CXCR3+ CM and CD28+/CD57+ EM, that exhibited significant changes during aging. Although CXCR3 is known as a marker for Th1 CD4 T cells and is required for migration to inflammatory sites^17–19^, the biological differences between CXCR3- and CXCR3+ CM subsets remain poorly characterized especially in CD8 T cells. To address this, we performed transcriptomic and epigenetic analyses to further characterize these two subsets. Since CD57 is known as a marker for senescent T cells, previously shown to increase with age^20–22^, we also included CD57+ EMRA in our analysis (Fig. 3A). Using a flow panel focused on these markers, we sorted these four memory CD8 T cell subsets as well as naïve CD8 T cells (Supplementary Table 1 and Supplementary Fig. 4) and performed both RNA-seq and CUT&RUN analyses. Principal component analysis (PCA) of transcriptomic data revealed distinct clustering of samples by subset, suggesting a unique transcriptomic profile in each (Fig. 3B). Differentially expressed gene (DEG) analysis across these CD8 T cell subsets identified 6,471 genes with significant differences (q-value < 0.05) (Fig. 3C). We then performed clustering to identify gene expression patterns across the CD8 T cell subsets, and identified seven gene sets with unique expression patterns (Fig. 3C). Among these, gene set 5 was specific to CXCR3- CM. Functional enrichment analysis of this gene set revealed associations with T helper cell differentiation and Th2 cytokine production (Figs. 3D, 3E and 3F, Supplementary Table 3), suggesting CXCR3- CM has a gene expression pattern resembling Th2 cells.

We then examined the transcript levels of individual T cell transcription factors, Th markers, effector molecules, and exhaustion markers across all subsets (Fig. 4A). As expected, we found that TCF7, LEF1, and SELL were high in naïve and both CM subsets, while TBX21 and Eomes were high in CXCR3+ CM, CD28+/CD57+ EM and CD57+ EMRA. Consistent with our functional enrichment analysis, GATA3 was high in CXCR3- CM, and GATA3-AS1, which promotes the expression of GATA3, was also elevated in this subset. Additionally, CXCR3 (a Th1 marker) was expressed in CXCR3+ CM, CD28+/CD57+ EM, and CD57+ EMRA, while CCR4, IL-4R, and PTGDR2 (Th2 markers) were expressed exclusively in CXCR3- CM. Surprisingly, CXCR3- CM did not express any of the cytotoxic effector molecules GZMA, GZMB, GZMK, GNLY, and PRF1, whereas CD28+/CD57+ EM expressed the highest levels of GZMK, suggesting that this subset resembles the GZMK+ age-associated T cells reported in previous studies^23^.

**Figure 4.**
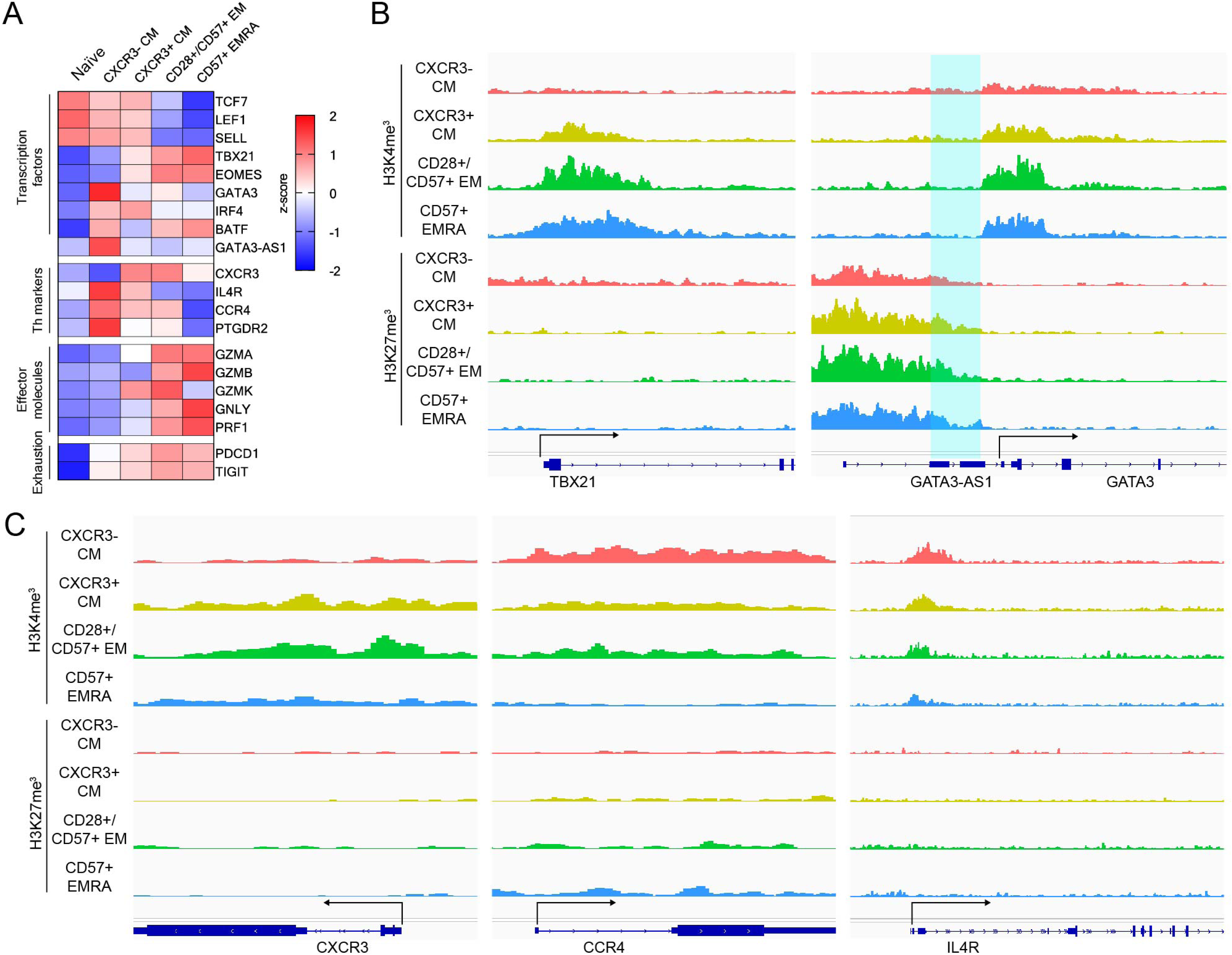
Epigenetic analysis exhibited unique epigenetic feature of CXCR3- CM T cells. **A.** Heatmap of gene expression of T cell transcription factors, Th markers, effector molecules and exhaustion markers in individual T cell subsets. **B.** IGV signal track of CUT & RUN with H3K4me3 (top) and H3K27me3 (bottom) at T cell transcription factors (Tbx21 (left) and GATA3 (right)) loci. Highlighted is the coding region of GATA3-AS1. **C.** IGV signal track of CUT & RUN with H3K4me3 (top) and H3K27me3 (bottom) at Th markers (CXCR3 (left), CCR4 (middle) and IL-4R (right)) loci.

These findings indicate that CXCR3- CM, which accumulate with age, exhibit a pattern of transcription characteristic of Th2 cells. This finding is supported by a recent scRNA-seq study of CD8 T cells from healthy human donors, which found that CD8 T cells expressing Th2 markers accumulate with age^13^. Our findings suggested that the CXCR3- CM population we identified using high-dimensional spectral flow cytometry is similar or identical to the cell population identified by Terekhova et. al., and spurred us to perform further experiments to better characterize this cell type and investigate possible mechanisms for its age-related accumulation.

### Histone methylation patterns in CXCR3- CM are distinct and compatible with Th2 T cells

To determine whether CXCR3- CM are epigenetically programmed to exhibit Th2 memory T cells transcriptional features, we analyzed their epigenetic landscape by using CUT & RUN on H3K4me3 and H3K27me3 marks. H3K4me3 is known as an active histone mark, whereas H3K27me3 is a repressive histone mark of transcriptional regulation^24,25^. We first examined the epigenetic status at genes of key T cell transcription factors (TCF7, LEF1, SELL, TBX21, EOMES and GATA3). Consistent with the RNA expression data, both CXCR3- CM and CXCR3+ CM exhibited high H3K4me3 signal at the TCF7, LEF1 and SELL loci and CD28+/CD57+ EM while CD57+ TEMRA showed elevated H3K27me3 signal at the LEF1 locus (Supplementary Fig. 5A). Intriguingly, CXCR3+ CM, CD28+/CD57+ EM and CD57+ TEMRA displayed high H3K4me3 and low H3K27me3 at the TBX21 locus. Conversely, CXCR3- CM displayed the opposite pattern, with low H3K4me3 and high H3K27me3 at TBX21 locus (Fig. 4B). Similarly, CXCR3- and CXCR3+ CM exhibited low H3K4me3 and high H3K27me3 at the EOMES locus, whereas CD28+/CD57+ EM and CD57+ EMRA exhibited high H3K4me3 and low H3K27me at this locus (Supplementary Fig. 4A). While the H3K4me3 signal at the GATA3 locus did not align with RNA expression, the H3K27me3 signal at GATA3-AS1, a non-coding RNA that promotes GATA3 expression, was higher in CXCR3+ CM, CD28+/CD57+ EM, and CD57+ TEMRA compared to CXCR3- CM (Fig. 4B). These findings demonstrated that CXCR3-CM cells have a distinct pattern of epigenetic features at T cell transcription factor loci.

Next, we examined the epigenetic features at the loci of Th subset-associated surface markers, (CXCR3, CCR4, IL4R, and PTGDR2). The H3K4me3 signal at CXCR3 was high in CXCR3+ TCM and CD28+/CD57+ EM, while the H3K27me3 signal did not show apparent differences between the subsets (Fig. 4C). In contrast to CXCR3, the H3K4me3 signal at Th2 maker loci (CCR4, IL4R, and PTGDR2) was more robust in CXCR3- CM compared to the other subsets (Fig. 4C), further supporting the idea that CXCR3- TCM carry distinct epigenetic features, aligning with their RNA expression profile. Finally, we analyzed the epigenetic features at the cytotoxic effector molecule loci GZMA, GZMB, GZMK, PRF1, and GNLY. CD28+/CD57+ EM and CD57+ TEMRA cells exhibited high H3K4me3 signals at GZMA and PRF1 loci (Supplementary Figs. 5B and 5C). Interestingly, CD28+/CD57+ EM showed a pronounced H3K4me3 signal at the GZMK locus, while CD57+ TEMRA exhibited a high H3K4me3 signal at the GZMB locus (Supplementary Fig. 5B), consistent with RNA expression in these subsets. Moreover, the H3K4me3 signal at GNLY was particularly high in CD57+ EMRA (Supplementary Fig. 5C). Taken together, these results indicate that each CD8 T cell memory subset has a distinct epigenetic signature at the loci of T cell transcription factors, Th subset markers, and effector molecules. Specifically, the epigenetic profile of CXCR3- CM is compatible with that of Th2 T cells.

**Figure 5.**
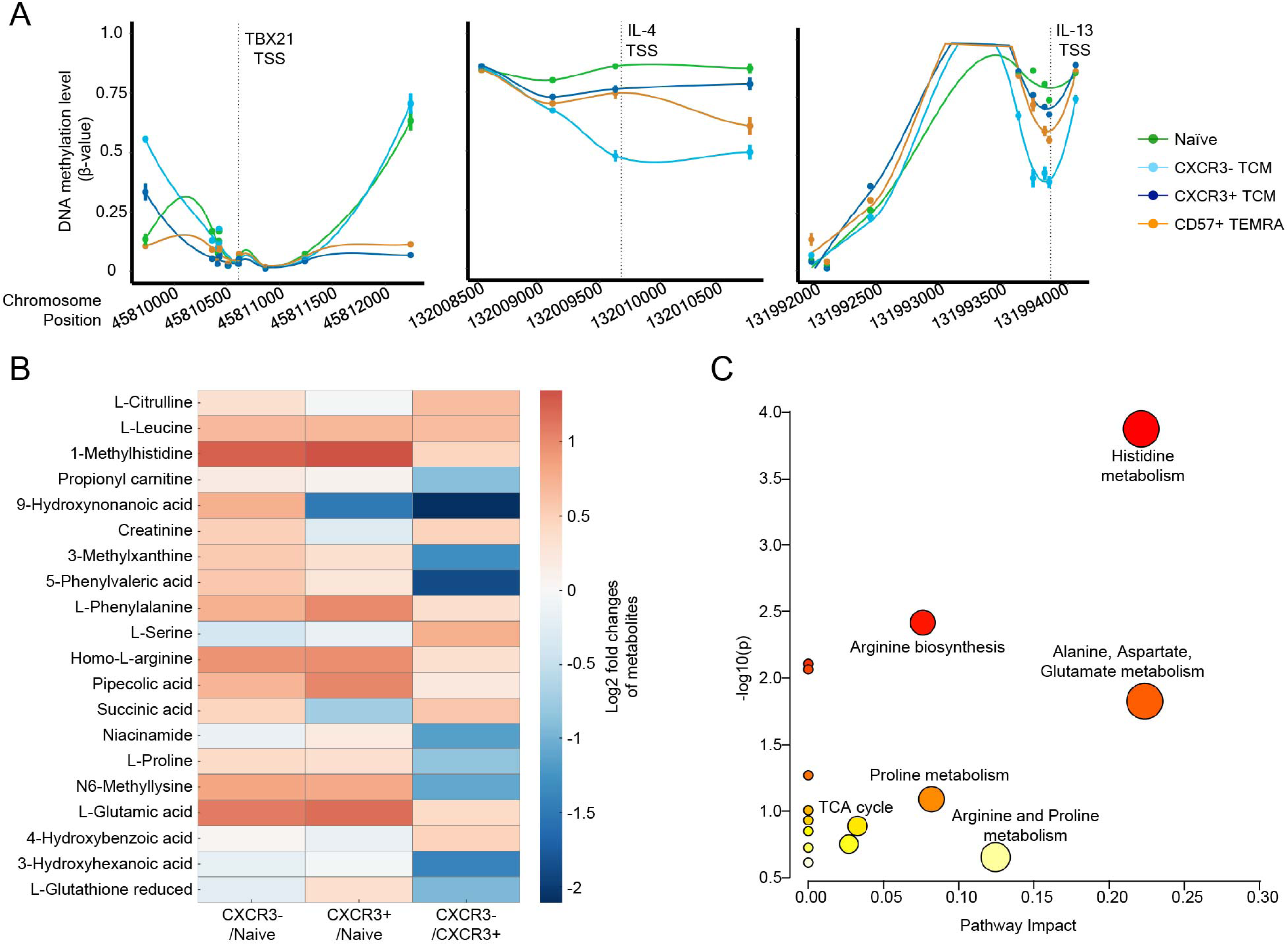
Additional multi-omics characterizations identified distinct DNA methylation and metabolomics signatures in CXCR3- TCM. **A.** DNA methylation level of CpG sites within 2 kbps of TBX21 (left), IL-4 (middle) and IL-13 (right) TSS in Naïve (green), CXCR3- TCM (light blue), CXCR3+ TCM (dark blue) and CD57+ EMRA (orange) are shown. **B.** Heatmap displaying the top differentially regulated metabolites across the three comparisons (CXCR3-/Naïve, CXCR3+/Naïve, and CXCR3-/CXCR3+) with clustering to show related patterns of change. Data were median-normalized, autoscaled, and analyzed using a paired t-test. Criteria for significance included a fold change ≥ 1.5 and a p-value ≤ 0.05. Red indicates upregulated metabolites, and blue indicates downregulated metabolites in each comparison. **C.** Pathway enrichment analysis comparing CXCR3- TCM CXCR3+ TCM. The plot highlights KEGG pathway analysis of significantly changed metabolites between the two subsets. Pathway significance is represented on the y-axis (−log10 p-value), while the pathway impact score is shown on the x-axis. Larger and darker points indicate pathways with higher levels of significance and impact, respectively.

### CXCR3- CM have a unique DNA methylation signature compared to CXCR3+ CM

To further explore epigenetic differences between CXCR3- and CXCR3+ CM, we performed DNA methylation analysis for these two subsets using DNA methylation arrays. Based on our histone modification findings, we focused on examining the DNA methylation patterns at loci associated with T cell transcription factors (TBX21 and GATA3), Th markers (CCR4 and IL4R) and Th2 cytokines (IL-4 and IL-13). In agreement with our transcriptomic and epigenetic data, CXCR3-CM cells exhibited distinct methylation patterns at loci associated with Th signature and immune response. Specifically, the DNA methylation level of CXCR3- CM at TBX21 was comparable to that of Naïve and higher than those of CXCR3+ CM and CD57+ EMRA (Fig. 5A). Although we did not observe apparent differences in DNA methylation levels between CXCR3- and CXCR3+ CM at GATA3, IL-4R and CCR4 (Supplementary Fig. 6), suggesting that histone modifications may play a more dominant role in the regulation of these genes, we did find that the Th2 cytokine loci (IL-4 and IL-13) were less methylated in CXCR3- compared to CXCR3+ CM (Fig. 5A). The expression of TBX21 in CXCR3- CM was thus suppressed by both DNA methylation and histone modifications. Furthermore, our findings suggest that the surface Th subset markers were primarily regulated by histone modifications, but Th associated cytokines were influenced by DNA methylation. These findings establish that CXCR3- CM not only exhibits a unique epigenetic and transcriptional profile consistent with Th2 memory T cells but also undergoes specific DNA methylation changes, highlighting their potential functional specialization during aging.

### Metabolic Adaptations of CXCR3- CM Favor Memory Formation

We performed metabolomics analysis on naïve, CXCR3-, and CXCR3+ CM subsets to explore whether these populations exhibit distinct metabolic profiles. Untargeted metabolomics on sorted CD8 subsets revealed separation between naïve and both types of CM, with partial overlap between the CXCR3- and CXCR3+ CM, as shown by supervised clustering analysis (Supplementary Fig. 7B). Of the 200 metabolites identified in these samples, significant differences between the subsets are shown in Fig. 5B. Upregulation of amino acid metabolism and nucleotide biosynthesis, crucial for supporting T cell proliferation and effector functions, were observed as expected in both CXCR3- and CXCR3+ CM compared to naïve, supporting activation and functional polarization^26^. CXCR3+ CD8+ T cells exhibit a reprogrammed metabolic state consistent with an effector phenotype, characterized by elevated levels of glycolytic intermediates, amino acid catabolites, and nucleotide biosynthesis products that support proliferation, cytotoxic activity, and inflammatory cytokine production. In contrast, CXCR3- CD8+ T cells display a more restrained metabolic signature, suggesting a memory state. The associated pathway enrichment analysis between the subsets highlights differential amino acid metabolism and TCA cycle intermediates (Fig. 5C). These metabolic shifts suggest distinct states of activation and differentiation between these subsets.

Interestingly, we also observed a significant decrease in short-chain fatty acids (SCFAs), isobutyric and isovaleric acids in CXCR3- CM compared to both naïve and CXCR3+ counterparts. These short-chain fatty acids promote T cell activation and function; thus, low levels of isobutyric acid in CXCR3- CM could reflect a less activated state^27^. Taken together, metabolic differences between CXCR3+ and CXCR3- CM reveal adaptations that align with their distinct immunological roles. The results suggest that CXCR3+ CM appears more predisposed to effector differentiation, whereas CXCR3- CM may be more inclined toward memory formation.

### CXCR3- CM cells express Th2 surface markers and produce Th2 cytokines upon stimulation

Our transcriptomic and epigenetic analyses revealed that CXCR3- CM exhibit key features of Th2 T cells. To further validate these findings, we modified our initial spectral flow panel to include the Th2 markers IL-4R, CCR4, and PTGDR2, and re-analyzed a subset of the samples, enabling further subclassification of CD8 T cell subsets (Supplementary Table 4 and Supplementary Fig. 7). We first examined the expression of each Th2 marker in the individual CD8 T cell subsets. Consistent with expectations, the expression level of IL-4R in CXCR3- CM was significantly higher than in Naïve and CD28+/CD57+ EM subsets. Similarly, the expression levels of CCR4 and PTGDR2 in CXCR3- CM were markedly elevated relative to other CD8 T cell subsets (Fig. 6A). These findings further confirm that CXCR3- CM exhibits distinctive features of Th2 T cells, a phenotype absent in other CD8 T cell subsets. The previous scRNA-seq study in healthy human donors identified CCR4 as a marker for Th2-like cells^13^. However, further analysis of Th2 markers within CXCR3- CM cells revealed additional complexity. As shown in Fig. 6B, CCR4 alone was not sufficient to define Th2-like cells. Specifically, 23.8% of CXCR3- CM cells were PTGDR2 single-positive, and 4% were IL-4R single-positive (Figs. 6C and 6D). These results suggest that CCR4 is expressed only in a subset of Th2 T cells and that the absence of CXCR3 expression may serve as a more reliable marker for identifying Th2-like T cells.

**Figure 6.**
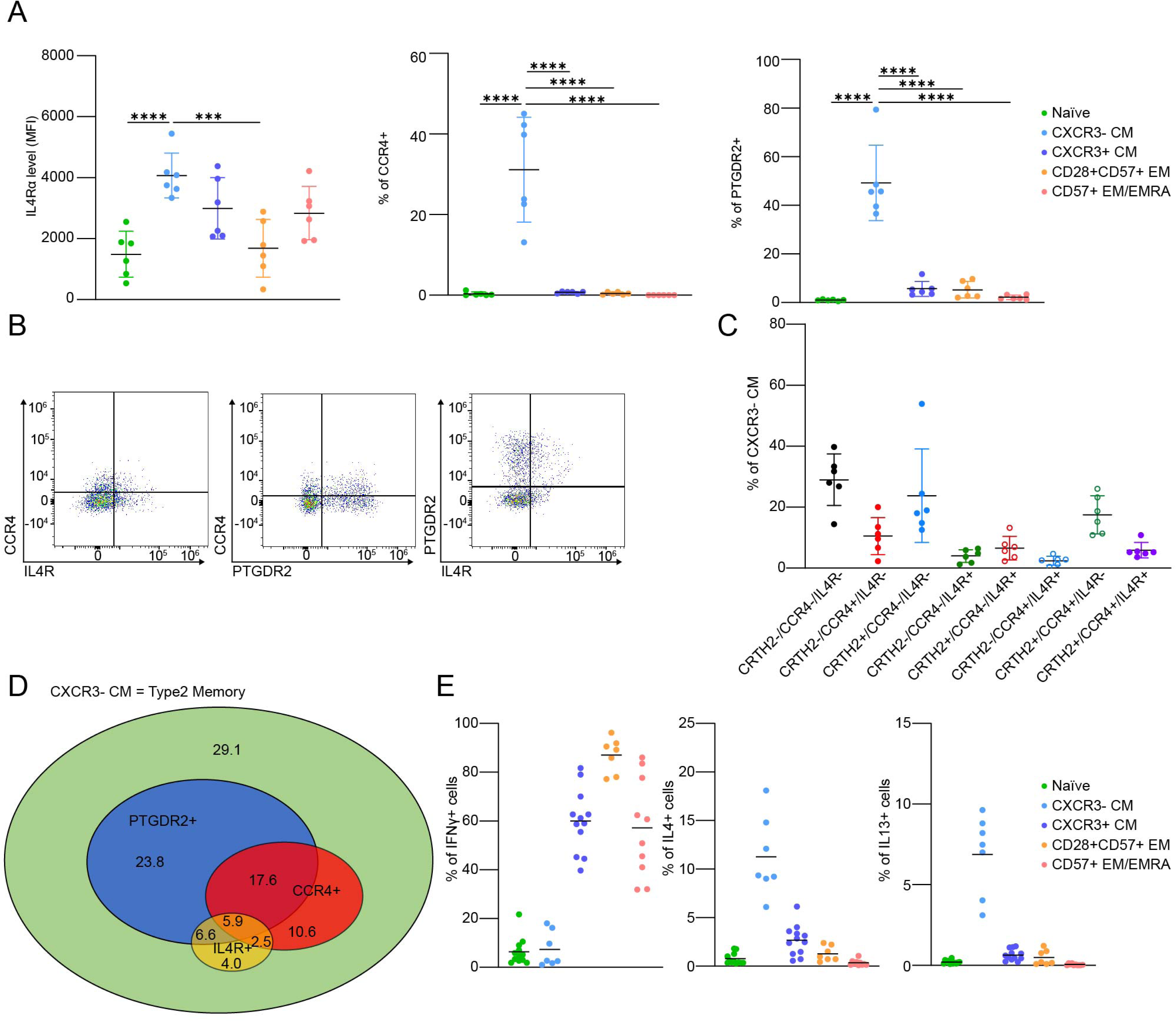
Flow cytometry analysis confirmed the expression of Th2 markers and the production of Th2 cytokines (IL-4 and IL-13) in response to PMA/Ionomycin stimulation in CXCR3- CM T cell subset. **A.** The expression levels of Th2 markers (IL-4R (left), CCR4 (middle) and PTGDR2 (right)) in individual T cell subsets are shown (n = 6, mean ± SD), One-way ANOVA; *** p < 0.001, **** p < 0.0001). **B.** Representative FACS plots of two of three Th2 markers in CXCR3- CM T cell subset (CCR4 vs IL-4R (left), CCR4 vs PTGDR2 (middle) and IL4R vs PTGDR2 (right)). **C.** Expression patterns of Th2 markers in CXCR3- CM T cell subset (n = 6, mean ± SD). **D.** Venn diagram of expression patterns of TH2 markers in CXCR3- CM T cell subset. Number shows the average percentage of individual expression patterns in CXCR3- CM T cell subset. **E.** The frequencies of IFN-γ, IL-4 and IL-13 producing cells in individual T cell subsets upon stimulation with PMA/Ionomycin (n = 7-12, mean ± SD, One-way ANOVA; **** p < 0.0001).

A previous study reported an accumulation of Th2 cells during aging and also that unfractionated CD8 T cells from old donors produce more Th2 cytokines than those from young donors^13^. However, it remains unclear whether CXCR3- CM specifically contribute to Th2 cytokines production. To investigate this, we measured the production of Th1 and Th2 cytokines across CD8 T cell subsets. We sorted isolated CD8 T cells into distinct subsets, stimulated them with PMA/Ionomycin *in vitro,* and analyzed them via flow cytometry using intracellular cytokine staining. Notably, far fewer CXCR3- TCM cells produced the Th1 cytokine IFN-γ in response to PMA/Ionomycin stimulation compared to the other CD8 T memory subsets (CXCR3+ CM, CD28+/CD57+ EM, and CD57+ EMRA). In accordance with our other findings, we confirmed that CXCR3- CM produced significantly higher levels of IL-4 and IL-13 upon stimulation than all the other subsets (naïve, CXCR3+ CM, CD28+/CD57+ EM and CD57+ EMRA). Importantly, none of the other CD8 T cell subsets produced IL-13, underscoring the unique functional capacity of CXCR3- CM cells (Fig. 6E). This cytokine profile suggests their functional specialization as Th2-like CD8 T cells.

### Naïve CD8 T cells from older donors are primed to make Th2 cytokines and show reduced expression of factors suppressing Th2 differentiation

Naïve CD4 T cells differentiate into Th2 CD4 T cells in the presence of IL-4, and central memory cells that are formed under these conditions are thereafter primed to adopt a Th2 effector phenotype^28^. CD8 T cells have also been shown to produce type 2 cytokines in response to specific stimuli^29^, suggesting that the CXCR3- CD8 CM we have observed may arise in a similar fashion after having been primed *in trans* as unbiased naïve cells. Alternately, some age-dependent change in the state of long-lived, mostly quiescent naïve CD8 T cells could result in a cell-intrinsic bias towards Th2-like responses. To test this hypothesis, we isolated naïve CD8 T cells from young and old donors, simulated them, and cultured them for one week in the presence or absence of IL-4 and anti-IFN-γ antibody (Non-polarizing vs Th2 conditions), then re-stimulated them briefly to examine the proportion of IFN-γ producing and IL-13 producing cells (Fig. 7A). About 25% of cells from young donors also did so under non-polarizing conditions and ∼17% under Th2 conditions, we found that naïve CD8 T cells from the older donors were somewhat (∼1.8-fold) more likely to produce IFN-γ than those from young donors (Fig. 7B). In contrast, very few cells from young donors made IL-13 in either non-polarizing or Th2 conditions (1-2%), while 6-8% of naïve cells from older donors did so (Figs. 7C, D). The ability to make type 2 cytokine responses therefore appears to be a property acquired by naïve CD8 T cells with age rather than the result of activated memory-forming cells being primed *in trans* by Th2 cytokines.

**Figure 7.**
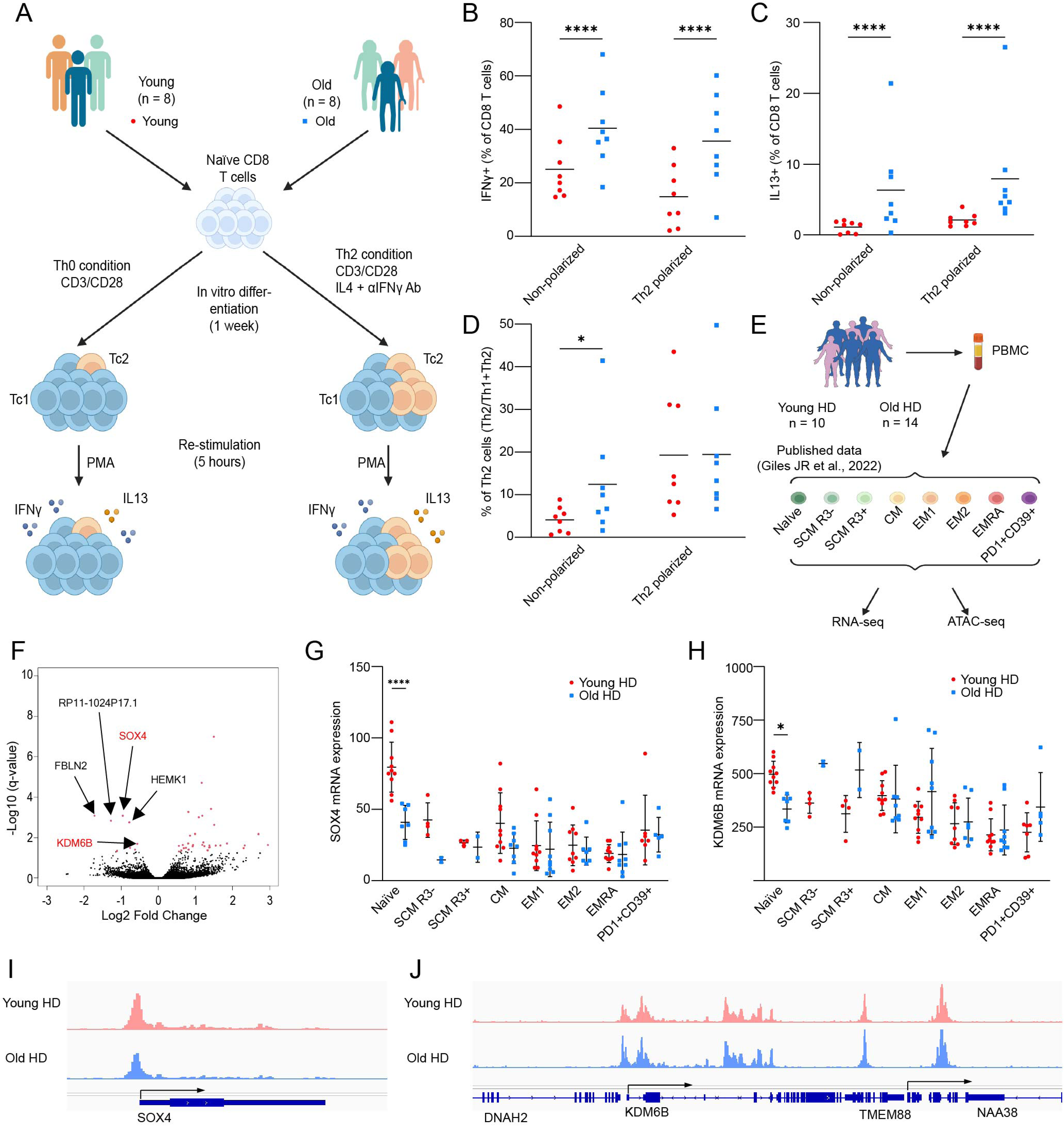
Old naïve CD8 T cells are more likely to differentiate into Th2 type memory T cells. **A.** Schematic workflow of in vitro CD8 Naïve T cell differentiation. **B-C.** The absolute frequencies of IFN-γ producing cells (B) and IL-13 producing cells (**C**) in individual conditions (n = 8, mean, Two way ANOVA; **** p < 0.0001). **D.** The proportion of Th2 cells (IL-13 producing cells) among both Th1 cells (IFN-γ producing cells) and Th2 cells in individual conditions. **E.** The schematic outline of the generation of public data set published in 2022. **F.** Volcano plot of the differentially expressed genes between young and old Naïve CD8 T cells. The genes with significance (q < 0.05) are shown in red dots. **G.** RNA expression of SOX4 across CD8 T cell subsets between young (red) and old (bule) healthy donors. **H.** RNA expression of KDM6B across CD8 T cell subsets between young (red) and old (bule) healthy donors. **I.** ATAC-seq signal track of young (red) and old (blue) healthy donors at SOX4 locus. J. ATAC-seq signal track of young (red) and old (blue) healthy donors at KDM6B locus.

To further investigate what changes in naïve CD8 T cells during aging might result in a bias towards type 2 effector function, we examined previously published bulk RNA-seq and ATAC-seq data from naïve CD8 T cells sorted from young and old, healthy donors^30^(Fig. 7E). Our DEG analysis on these data identified 33 genes significantly increased in older donors vs. younger ones, and 8 genes significantly decreased. Among the significantly decreased genes, we identified the transcription factor SOX4 and the histone demethylase KDM6B (Fig. 7F). only in naïve CD8 T cells (Figs. 7G and 7H). The ATAC-seq data also showed increased accessibility at the SOX4 locus in young donors vs. old donors, providing further evidence of its reduced expression in older naïve CD8 T cells (Figs. 7I and 7J). Importantly, both of these genes have known functions in the regulation of T cell differentiation: SOX4 suppresses GATA3 function by interfering with its binding to DNA, thus inhibiting Th2 differentiation^31^, and KDM6B supports Th1 differentiation by demethylating H3K27me3 at TBX21 locus^32^. Taken together, these findings provide a possible mechanistic explanation for the age-dependent increase in Th2-like CD8 T cells that we have observed.

### The frequency of CXCR3- CM cells is a biomarker for asthma, type 2 diabetes and chronic liver disease

Type 2 immunity plays an important role in the pathogenesis of allergic diseases, including asthma and atopic dermatitis, however, it remains unknown whether the accumulation of CXCR3- TCM has any impact on health outcomes. To investigate this, we performed health outcome logistic regression analysis based on DNA methylation (DNAm) patterns. Specifically, we first flow-sorted CXCR3- and CXCR3+ CM, extracted DNA from 10 donors and acquired DNAm profiles using the EPIC v2 array. We then merged these data with DNAm data from a previous study^33^, to construct a novel DNAm reference matrix encompassing our sorted cells and 5 other leucocyte subtypes from their data (monocytes, neutrophils, CD4 T-cells, B cells, and NK cells). We used this matrix to impute cell frequencies from whole blood DNA methylation datasets in the Massachusetts General Brigham Aging Biobank Cohort (MGB-ABC), which consists of 4,386 donors, and provides comprehensive electronic medical records of the participants (Fig. 8A). We then performed logistic regression analysis to evaluate the association between each immune cell fraction and health outcomes (Fig. 8A). In addition to showing the known positive associations between the frequency of monocytes and cardiovascular diseases (Odds ratio (OR): 1.16, 95% Confidence interval (CI): 1.08-1.25)^34^ (Fig. 8B), our analysis showed a positive association between the estimated frequency of CXCR3-CM cells and asthma (OR: 1.12, 95% CI: 1.05-1.19), suggesting that CXCR3- CM can mediate pathologies associated with Th2 responses. Unexpectedly, the frequency of CXCR3- CM also had a positive association with multiple chronic liver diseases (OR: 1.17, 95% CI: 1.09-1.26), as well as type 2 diabetes (OR: 1.11, 95% CI: 1.04-1.19) (Fig. 8B and 8C). Several of these diseases do not have clear autoimmune etiologies, however type 2 inflammation is known to contribute to liver fibrosis, so our findings may indicate a previously unappreciated role for CXCR3- CM-mediated autoinflammation in these conditions^35^. Th2-driven inflammation observed in CXCR3- CM may contribute to fibrotic complications associated with diabetes and liver disease. Elevated Th2 cytokines, including IL-4 and IL-13, have been linked to increased fibrosis in both adipose and liver tissues, potentially exacerbating insulin resistance and liver fibrosis. Additionally, intrinsic changes in naïve CD8 T cells towards Th2 polarization and subsequent accumulation of CXCR3- CM during aging, along with its correlation with these diseases, suggest possible underlying mechanisms. Our data show that the accumulation of CXCR3- CM, characterized by a distinct epigenetic, transcriptomic and metabolic profile compatible with Th2-like T cells, contributes to Th2 inflammation, which manifests in accelerated aging and aging-related complications.

**Figure 8.**
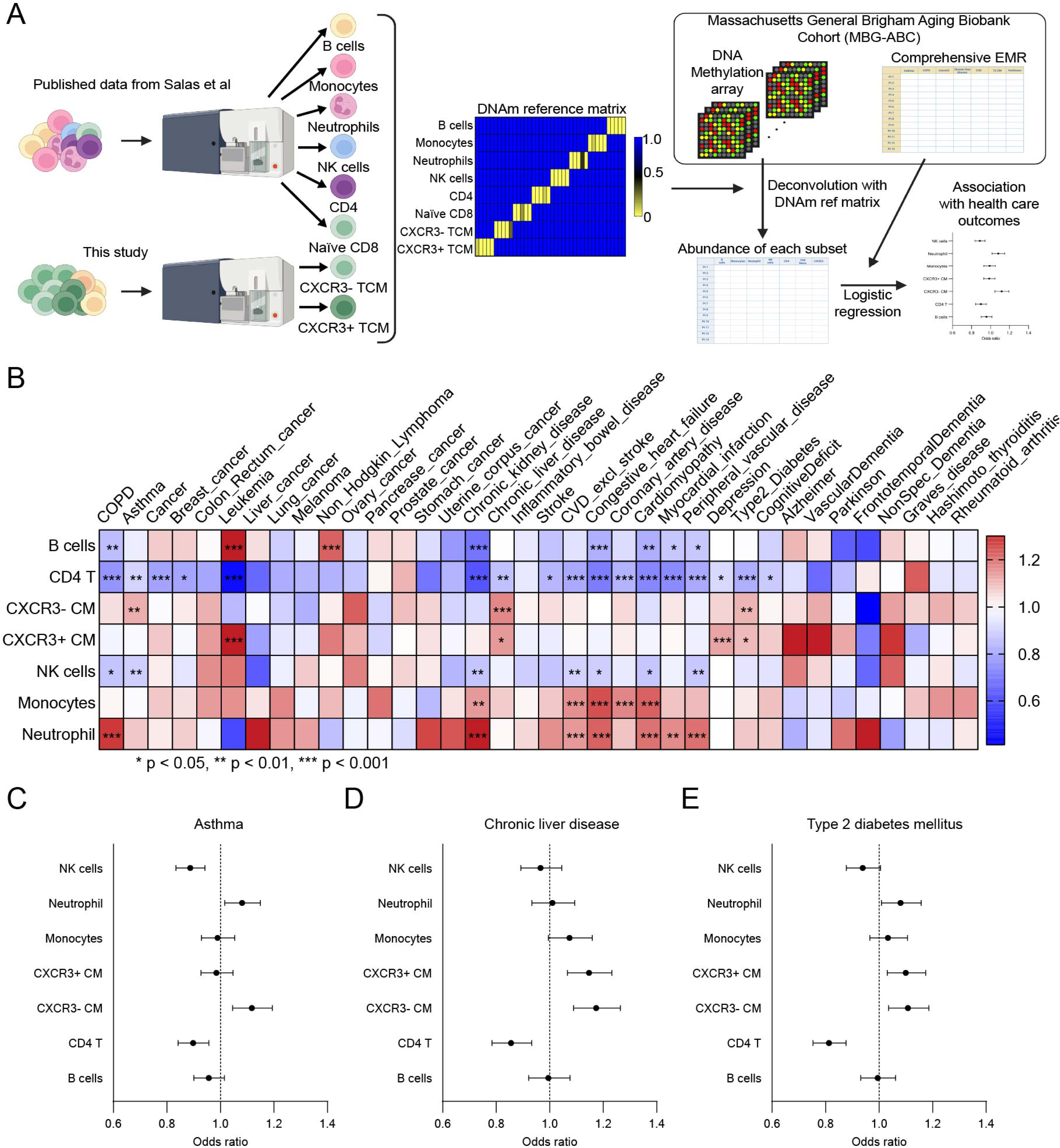
The abundance of CXCR3- CM is a potential biomarker for asthma, chronic liver disease and type 2 diabetes mellitus. **A.** Schematic workflow for evaluating associations of immune cell fractions with health care outcomes using MBG-ABC. **B.** The odds ratios of the associations between each immune cell fraction with health care outcomes were shown in the heatmap (Logistic regression, * q < 0.05, ** q < 0.01, *** q < 0.001). **C-E.** Forest plots of associations between each immune cell fraction with asthma (**D**), chronic liver disease (**D**) and type 2 diabetes mellitus (**E**). Odds ratios are shown in black dots and error bars indicate 95% CIs.

## Discussion

In this study, we have employed spectral flow cytometry to analyze age-related changes in blood immune cell composition, employing a 36-color panel focused on markers of CD8 T cell differentiation and exhaustion. Of particular interest, we find that CXCR3- central memory CD8 T cells accumulate during aging, in a robust manner that is independent of CMV serostatus or sex. Their CXCR3+ counterparts also accumulate with age, but at a lower rate and from a higher baseline, resulting in a shift from the near absence of CXCR3- cells in young donors to an abundance comparable to that of CXCR3+ cells in most older donors.

Our subsequent transcriptomic, epigenetic, and functional analyses of this age-accumulating population have revealed that unlike CXCR3+ CM, which express T-bet, have activating epigenetic marks at Th1-associated loci, and elaborate cytotoxic effector programs, CXCR3- CM express GATA3, have activating epigenetic marks in Th2-associated loci, and present a more Th2-like effector program. Importantly, we have also found that with age, naïve CD8+ T cells become progressively more primed to make Th2-like responses upon stimulation, suggesting that cell-intrinsic changes in the naïve pool drive the emergence of CXCR3- CD8+ T cells rather than remodeling of the SCM/CM pool *in trans* during immune responses.

In support of this idea, transcriptomic analysis of naïve CD8 T cells from old vs young donors shows reduced expression of KDM6B, a histone demethylase essential for repressing the Th2 phenotype, as well as Sox4, a repressor of GATA3 function, suggesting a mechanism for this age-dependent bias^31,32^. Together, these findings support a model in which progressive, age-dependent degradation of the epigenetic programs of naïve CD8 T cells, responsible for tight regulation of their effector polarization in response to appropriate stimuli, results in the emergence of a potentially pathogenic population of aberrantly primed CD8 memory cells. Age-dependent changes in DNA methylation patterns are a well-established phenomenon and occur predictably enough that they have become the basis of several molecular clocks for measuring organismal age^36–39^. Moreover, the gradual corruption of the gene regulatory programs that tightly control proliferation and maintain cellular identity is a broadly recognized mechanism in the age-dependent increase in dysplasia and malignancy, and the emergence of mis-primed memory CD8 T cells could be an instance of this phenomenon occurring in the immune system. Better understanding the specific mechanism behind this age-dependent degradation of epigenetic information in naïve CD8 T cells will be a major topic in our further investigations going forward.

Our flow analysis has provided some additional insights into memory T cell composition changes during aging beyond the accumulation of Th2-like CXCR3- CM, also identifying CD28+/CD57+ EM as an age-accumulating subset. A recent study in mice identified age-associated CD8 T cells characterized by their co-expression of a specific set of markers (GZMK, PD-1, CD49d, TOX, and EOMES)^23^. Our own transcriptomic and epigenetic analysis revealed that CD28+/CD57+ EM cells express GZMK but not GZMB, thus resembling these age-associated CD8 T cells. A recently published scRNA-seq atlas of healthy human PBMCs identifies CCR4 as a potential defining surface marker of Th2-like CD8 T cells^13^ Our own surface marker analysis suggests that CCR4+ CD8 CM only constitute a subset of Th2-like CD8 T cells, and that loss of CXCR3 expression tracks more closely with this cellular phenotype.

An important unanswered question is whether the accumulation of CXCR3- CM with age has any impact on health. Our logistical regression analysis based on whole blood DNA methylation patterns in a large aging-focused biobank (MGB-ABC), indicates a positive association between the abundance of CXCR3- CM and several chronic diseases, including asthma, type 2 diabetes, and both autoimmune and non-autoimmune liver disease.

Type 2 responses are required for effective tissue repair, but if prolonged and/or excessive, they can lead to fibrosis^35,40^. Prior work in animal models shows that IFN-γ deficient mice develop non-alcoholic steatohepatitis (NASH) with fibrosis, while obese IL-4/Il-10 knockout mice are protected against NASH^41^. Furthermore, *Stat1* -/- mice, which lack the ability to mount Th1 responses, exhibit Th2-mediated biliary duct injury after infection, leading to biliary atresia^42^. These findings highlight the importance of tight regulation of the Th1/Th2 balance to maintain appropriate immune responses and establish a clear connection between Th2-favoring imbalances and chronic liver disease. Importantly, these findings also suggest that the accumulation of CXCR3- CM could serve as a readily assayed blood biomarker for liver fibrosis. Expanding this investigation to include functional assays demonstrating the pathogenic capacity of CXCR3- CM in liver fibrosis models could solidify this link and provide a basis for therapeutic intervention.

In conclusion, our study highlights the accumulation of CXCR3- CM with Th2 phenotypes as a novel hallmark of aging immune systems. These cells likely arise due to intrinsic epigenetic changes and may contribute to chronic inflammatory conditions, such as liver fibrosis. Future studies focusing on functional characterization and comparisons with other T cell subsets would shed further new light to deepen our understanding of aging-related immune dysregulation.

## Limitations

This study has some limitations that should be addressed in future research. First, age group stratification was not performed due to the wide range of age distribution among donors.

Stratifying age groups could provide a more detailed understanding of progressive aging phenotypes in the context of cellular subset changes. Additionally, while we observed a significant decrease in KDM6B expression in CD8+ T cells from older donors, we were unable to demonstrate that the KDM6B locus itself undergoes epigenetic changes during aging. This suggests that other regulatory factors may contribute to the reduced expression of KDM6B, highlighting an important potential avenue for future investigation.

## Methods

### Human Blood donor samples

Leukapheresis products from healthy blood donors are purchased from Vitalant® (San Francisco, CA, USA). Donor information regarding age, sex, and CMV serostatus were provided by Vitalant®. We collected the human PBMC from 45 blood donors with an almost equal number of CMV negative and CMV positive donors as well as female and male donors. Peripheral blood mononuclear cells were purified from them by Ficol-paque density gradient centrifugation. CD8 T cells were directly isolated from them by using RosetteSep^TM^ human CD8+ T cell enrichment cocktail (STEMCELL^TM^ TECHNOLOGIES).

### Cell culture

Isolated PBMC and CD8 T cells were cultured in complete RPMI (RPMI1640 with 10% FBS (Gemini), penicillin and streptomycin (100IU and 100ug/ml, Corning), 1mM sodium pyruvate (Corning), HEPES (100x, Cytiva HyClone, Fisher) and 50 IU/ml human recombinant IL-2 (PeproTech).

### CyTEK Aurora, surface staining

Cryopreserved PBMCs were thawed and cultured in cRPMI overnight. After cell counting, 2 million PBMCs were subjected to staining. Cells were resuspended in RPMI with 10% FBS and 1uM bafilomycin and incubated at 37°C for 1.5 hours. After incubation, Spider-β-Gal (Dojindo) was added to the samples at a concentration of 20μM and incubated for 30 minutes. Then, cells were stained with LIVE/DEAD Fixable Blue dead cell stain kit (Invitrogen, L23105) at 4°C for 30 minutes. After washing, cells were resuspended in FACS buffer with human IgG (Sigma-Aldrich, I4506) for Fc block and incubated for 30 minutes. After Fc blocking, an antibody cocktail was added to the samples and samples were incubated for an hour. After washing 2 times with PBS, cells were resuspended in FACS buffer and acquired on Aurora® spectral flow cytometer. The list of antibodies used were provided in the Supplementary Table 1(Initial panel) and 4(Th2 panel).

### CyTEK Aurora, intracellular staining

Stimulated CD8 T cells were stained with LIVE/DEAD Fixable Blue dead cell stain kit at 4°C for 30 minutes, followed by Fc blocking with human IgG for 15 minutes and staining with a surface antibody cocktail for 30 minutes at 4°C. Then, fixation and permeabilization were performed for 15 minutes at RT using Foxp3/Transcription Factor Fixation/Permeabilization Concentrate and Diluent kit (eBioscience, #00-5521-00). After permeabilization, samples were stained with an intracellular antibody cocktail for 30 minutes at RT and run on Aurora®.

### Spectral flow data analysis

Data were unmixed, further manually compensated and exported as fcs files using SpectroFlo® software. Exported fcs files were imported into FlowJo and samples were manually gated to exclude cell debris, doublets, dead cells and doublets co-expressing CD3/CD14, CD14/CD19, CD14/CD56, CD3/CD19 and CD19/CD56. Next, CD45+ leukocytes were manually gated and used for downstream analysis. We subsampled CD45+ leukocytes to 20,000 events per donor using DownSample v3 plugin to both reduce cell numbers and accommodate the different number of cells resulting from doublet removal. The subsampled CD45+ leukocytes were concatenated into a fcs file to perform dimensionality reduction analysis. Dimensionality reduction analysis was performed using the UMAP plugin on the FlowJo. CD8 T cells were further subsampled to 12,000 events per donor and concatenated into a fcs file to perform downstream analysis. Dimensionality reduction analysis of CD8 T cells was performed using the UMAP plugin on the FlowJo. Clustering was performed using FlowSOM plugin on the FlowJo with following markers; CD45RA, CCR7, CD95, CD27, CD28, CD49d, CD127, CD2, CXCR4, CD25, HLA-DR, CD38, CCR6, CXCR3, CD39, PD-1, KLRG1, TIGIT, SLAMF4, CD57, CD16, CD56, NKG2A, NKG2D, CD161 and KIR.

### FACS sorting

Freshly isolated CD8 T cells were cultured at 37°C overnight in cRPMI. Cells were washed and stained with LIVE/DEAD Fixable Near-IR dead cell stain kit (Invitrogen, L10119) at 4°C, followed by Fc blocking with human IgG (SIGMA, I4506) for 15 minutes at 4°C and staining with an antibody cocktail for one hour at RT. Samples were sorted on FACS Aria II machine into five CD8 T cell subsets. A small aliquot of sorted samples was run to check purity. Sorted cells were washed once with PBS and used for downstream assays. The list of antibodies for sorting is provided in the Supplementary Table 2.

### In vitro T cell stimulation assay

FACS sorted cells were cultured at 37°C overnight. Cells were stimulated with 15 ng/ml of PMA and 500nM of ionomycin for 5 hours in the presence of 5 μg/ml of Brefeldin A (Biolegend). After stimulation, samples were stained and run on CYTEK Aurora as described above.

### In vitro T cell differentiation assay

Cryopreserved PBMCs were thawed, resuspended in cRPMI with 50 IU/ml human recombinant IL-2 and cultured at 37°C overnight. Naïve CD8 T cells were isolated from PBMCs using EasySep Human Naïve CD8+ T Cell Isolation Kit (STEMCELL TECHNOLOGIES, #19528). Isolated Naïve CD8 T cells were stimulated with plate-bound anti-CD3 (2.5 μg/ml, Tonbo Bioscience, clone UCHT1) and soluble CD28 (5μg/ml, Tonbo Bioscience, clone CD28.2) in cRPMI with 50 IU/ml human IL-2 in the absence (Non-polarizing) or presence (Th2-polarizing) of anti-IFN-γ (5 μg/ml, Biolegend, 506532) and human recombinant IL-4 (10 ng/ml, Proteintech, HZ-1004) in 24-well plate for 3 days with a medium on day 2. After 3 days of stimulation, samples were transferred into new 12-well plates to stop stimulation and cultured at 37 for another 4 days in cRPMI with 50 IU/ml IL-2 in the absence or presence of IFN-γ and IL-4. After 4 days of culture, samples were re-stimulated with PMA and ionomycin for 5 hours and stained with surface and intracellular antibodies as described above. These samples were run on Aurora.

### DNA and RNA-extraction

Genomic DNA was extracted from the FACS sorted CD8 T cells using Quick-DNA Microprep kit (Zymo Research, D3020) per manufacturer’s protocol. RNA was extracted from the FACS sorted CD8 T cells using Quick RNA Microprep kit (Zymo Research, R1050) per manufacturer’s protocol.

### Sample preparation for CUT & RUN

Of the FACS sorted CD8 T cells, 100,000 from each sorted population was collected for each respective immunoprecipitation pull down assay and an input sample. Some modifications were made to the manufacturer’s protocol for the CUT & RUN Assay Kit (Cell Signaling Technologies, #86652). Briefly, live cells were washed (+spermine, +PIC) and mixed with activated Concanavalin A beads. After incubation, for IP samples: antibody binding buffer (+spermidine +PIC +digitonin) was added to clean cell:bead suspensions. Each reaction was incubated overnight with the respective antibody; anti-H3K4me3 (CST-C42D8), anti-H3K27me3 (CST-C36B11), anti-Rabbit IgG (CST-DA1E). Cells were washed and lysed with Digitonin Buffer (+spermidine, +PIC, +digitonin) and resuspended in the presence of pAG-MNase. Excess pAG-MNase was washed, and Calcium Chloride was added to activate the enzyme. The reaction was stopped with Stop Buffer (+digitonin, +RNase A, +spike-in DNA), and spun to collect the supernatant containing the fragmented DNA. For input samples: cells were incubated and lysed in successive steps with Buffers A & B (CST#14282). Diluted MNase (CST#10011) was added to fragment the DNA, digestion was stopped using EDTA before collecting DNA in the supernatant. Proteins and RNAs were digested with the addition of proteinase K and RNAse A. Fragmented DNA was purified using Spin Columns (CST#14209) per manufacturer’s protocol.

### RNA-seq data processing and analysis

Quality control and trimming of original FASTQ files were performed using Rfastp package with default parameters. QC and trimmed FASTQ files were aligned using subjunc function of Rsubread package with the hg38 human reference genome. Generated BAM files were sorted and indexed using Rsamtools. Transcript counts per gene were generated using the summarizeOverlaps function of GenomicAlignments package. Normalization and log transformation of gene counts were performed using the rlog function of DESeq2 package. PCA was done with the plotPCA function of DESeq2 package using all the genes. Differentially expressed genes (DEGs) across CD8 T cell subsets were identified by running likelihood-ratio test with DESeq2 package using adjusted p value < 0.05.

### Clustering

K-means clustering was performed using the pheatmp function from pheatmap package to identify the groups of significant DEGs that have similar expression patterns across subsets.

### Enrichment analysis

Functional enrichment analysis was performed using STRING (https://string-db.org/) using all genes as background gene list.

### CUT & RUN Data processing

Quality control and trimming of original FASTQ files were performed using Rfastp package with default parameters. QC and trimmed FASTQ files were aligned using subjunc function of Rsubread package with the hg38 human reference genome. Generated BAM files were sorted and indexed using Rsamtools. Sorted and indexed BAM files were converted to Bigwig file by using coverage to create RLElist objects and then export.bw function in the rtracklayer to export them as bigwig files. Average Bigwig files were created by combining the bigwig files from all the donors in individual conditions with bigwigAverage.

### DNA methylation assessment

DNA extracted from FACS sorted CD8 T cell subsets was bisulfite-converted by using EZ DNA Methylation kit (Zymo Research) following manufacturer’s protocol. Bisulfite-converted DNA was assigned to the wells on the Infinium HumanMethylationEPIC BeadChip. After amplification, hybridization, staining and washing, raw image intensities were taken by using Illumina iScan SQ instrument and saved as IDAT files.

IDAT files were further converted into beta values by using the minfi R package^43^, with a functional normalization pre-processing step^44^. DNA methylation data from Naïve and CD57+ TEMRA were obtained from our previous study and included in the analysis after batch correction^45^. CpG sites located within 1500 bp upstream or downstream of the transcription start site were analyzed.

### Published RNA-seq and ATAC-seq data processing and analysis

Dataset for RNA-seq were downloaded from the Gene Expression Omnibus database (GEO) under GSE179609 with the normalized, log2 transformed and batch corrected expression tables. The RNA expression table was imported into R. Differentially expressed gene analysis between young and old healthy donors in individual CD8 T cell subsets were performed using the edgeR package. Dataset for ATAC-seq were downloaded from the GEO under GSE179613. Downloaded SRA files were converted to fast files using SRA tool. Quality control and trimming of original FAST files were performed using the fastp. QC and trimmed FASTQ files were aligned using HISAT2 with the hg38 human reference genome. Generated BAM files were sorted and indexed using samtools. Sorted and indexed BAM files were converted to Bigwig files by using bamCoverage in the deeptools package. Average Bigwig files were created by combining the bigwig files from all of the young or old donors with bigwigAverage.

### Sample Preparation for Metabolomics

Metabolites were extracted from snap-frozen cell pellets of CD8 T cell subsets using a biphasic extraction method adapted from Matyash et al. (2008) and Ding et al. (2021). This method utilized methanol (MeOH) and methyl tert-butyl ether (MTBE) to ensure efficient recovery of polar and non-polar metabolites. To initiate the extraction, 225 μL of methanol containing internal standards was added to the frozen cell pellets, followed by vortexing for 10 seconds to disrupt cellular material. Next, 750 μL of MTBE containing additional internal standards was introduced, and the samples were shaken for 10 minutes at 4 °C to enhance metabolite solubilization. Subsequently, 188 μL of ultrapure water was added to the samples to induce phase separation, followed by vortexing for another 10 seconds.

The samples were centrifuged at 14,000 × g for 2 minutes, yielding distinct organic (upper) and aqueous (lower) phases. These phases were carefully transferred into separate 1.5 mL tubes and dried using a speed vacuum concentrator. The extraction process provided a comprehensive preparation of metabolites suitable for downstream analyses.

### Chromatographic Separation

Chromatographic separation was performed on an ACQUITY UPLC CSH BEH column (1.7 μm, 100 × 2.1 mm; Waters Corporation, Milford, MA, USA) installed on a Vanquish Horizon UHPLC system (Thermo Fisher Scientific, Waltham, MA, USA). For untargeted metabolomics, the mobile phases consisted of Buffer A (water containing 10 mM ammonium formate and 0.125% formic acid) and Buffer B (95% acetonitrile and 5% water, containing 10 mM ammonium formate and 0.125% formic acid). A gradient elution method was employed, starting with a high proportion of Buffer A to elute polar metabolites, and gradually increasing the proportion of Buffer B for hydrophobic compounds. The flow rate was set at 0.5 mL/min, and the system was equilibrated between runs to ensure consistent retention times. To prevent sample carryover, the injection system was washed before and after each sample with a solution of 90:10 water/methanol containing 0.1% formic acid. This protocol ensured high reproducibility and optimal separation of metabolite species.

### Data Acquisition and Analysis

Data were acquired using an Orbitrap IQ-X Tribrid mass spectrometer (Thermo Fisher Scientific, Waltham, MA, USA) equipped with an EASY-Max NG ion source operating in positive ion mode. The electrospray ionization source parameters are listed in Supplementary Table 5.

Mass spectrometry was performed in data-dependent acquisition (DDA) mode. Full MS scans were collected in the Orbitrap analyzer, with real-time internal calibration using fluoranthene ions provided by the EASY-IC™ system. The intensity threshold for triggering fragmentation was set to 2.0 × 10⁴, and higher-energy collisional dissociation (HCD) was used to fragment selected ions. Dynamic exclusion was applied to minimize repeated fragmentation of abundant ions and maximize metabolite coverage. Data processing and peak annotation were conducted using Compound Discoverer (Thermo Scientific), with alignment performed against in-house and public metabolite databases. Statistical analyses and pathway enrichment studies were carried out using MetaboAnalyst and R packages enabling detailed interpretation of the metabolomic data.

### Construction of reference matrix

To estimate immune cell fractions from whole blood DNA methylation data, a reference matrix was constructed by integrating deconvoluted immune cell profiles with previously published methylation datasets. DNA methylation data from this dataset were combined with a previously generated dataset^33^ to create a reference matrix of DNA methylation patterns per cell type. The Houseman regression method was applied to generate cell-type-specific methylation signatures, which were integrated into a reference matrix for deconvolution. Any negative cell fraction values were replaced with zero to ensure biological plausibility, and all estimates were z-score transformed before further analysis. Cell type predictions were validated by predicting cell type proportions on individually sorted cells (Supplemental Fig. 9).

### Clinical Outcomes and Disease Classification in MGB-ABC Cohort

Disease outcomes were derived from electronic health record (EHR) data within the MGB-ABC cohort. Associations with prevalent diseases were evaluated with adjustment of age and sex in the logistic regression model. The chronic liver disease category included alcoholic liver disease, non-alcoholic fatty liver disease (NAFLD), chronic viral hepatitis, cirrhosis, and other chronic liver diseases.

Additional conditions analyzed included asthma, type 2 diabetes, chronic kidney disease, inflammatory bowel disease, and cardiovascular diseases (coronary artery disease, congestive heart failure, myocardial infarction, and peripheral vascular disease). Associations were also tested for multiple cancers, including breast, colorectal, leukemia, liver, lung, melanoma, non-Hodgkin lymphoma, ovarian, pancreatic, prostate, stomach, and uterine corpus cancers.

Neurological and psychiatric disorders were examined, including manifestation of cognitive deficits, Alzheimer’s disease, vascular dementia, frontotemporal dementia, non-specific dementia, and Parkinson’s disease, as well as autoimmune conditions such as Graves’ disease, Hashimoto’s thyroiditis, and rheumatoid arthritis. Clinical phenotypes were identified using standardized diagnosis codes and validated through EHR-based phenotype definitions.

### Statistical Analysis for Immune association to disease

Associations between immune cell fractions and clinical outcomes were assessed using cross-sectional logistic regression for disease prevalence risk.

For cross-sectional analyses, logistic regression models were fitted using the glm() function in R with a binomial link function to estimate the association between immune cell fractions and the presence of disease:

logit(P(Y=1))=β0+β1⋅Cell Fraction+β2⋅Age+β3⋅Sex+ɛ\text{logit}(P(Y=1)) = \beta_0 + \beta_1\cdot \text{Cell Fraction} + \beta_2 \cdot \text{Age} + \beta_3 \cdot \text{Sex} + \epsilonlogit(P(Y=1))=β0 +β1 ⋅Cell Fraction+β2 ⋅Age+β3 ⋅Sex+ɛ

where Y represents a binary disease outcome, Cell Fraction denotes the deconvoluted immune cell proportion, and age and sex were included as covariates.

All models were adjusted for age and sex, and results were reported as odds ratios (OR) with 95% confidence intervals (CIs) for logistic regression. Multiple comparisons were corrected using the Bonferroni method to control the false discovery rate.

## Acknowledgement

This research was funded by Hevolution Foundation to the Buck Institute for Research on Aging (HF-PART-23-1422047), National Institute on Aging Grants (T32 AG000266), National Institutes of Health (U01AI180158), and by institutional support from the

Buck Institute for Research on Aging.

## Contributions

H.M., M.C., P.V.A.K., B.D.A. designed, performed, and analyzed the experiments with help from R.T., R.K., R.R.R., S.I., A.F., G.V.H., C.A. and H.G.K. H.M. and M.C. designed flow panel with help from R.T., R.K. and H.G.K. A.T., V.D., Q.C., J.LS. performed health outcome logistic regression analysis with help from R.S. J.LG., G.V.H., R.R.R., H.G.K, and E.V. assisted in manuscript drafting. P.V.A.K., and B.D.A. performed the metabolomics experiments with help from B.S. J.LG. assisted in data analysis of Cut and Run experiments. H.M., M.C., M.M.K., drafted and wrote the manuscript. All authors contributed to manuscript editing and revising.

## Tables

**Supplementary Table 1. Spectral flow cytometry panel used in the analysis of human peripheral blood mononuclear cells.**

**Supplementary Table 2. Flow cytometry panel used for sorting of five CD8 T cell subsets. Supplementary Table 3. Enrichment analysis results in individual gene set.**

**Supplementary Table 4. Spectral flow cytometry panel used to validate the expression of Th2 markers.**

**Supplementary Table 5. The electrospray ionization source parameters used in the metabolomics analysis.**

## Conflict of Interest Statement

The authors declare that they have no conflict of interests.

## Data Availability Statement

The raw data supporting the findings of this study will be openly available in various repositories and all processed data will be available as supplementary materials of the article. Raw flow cytometry data will be accessible on the NIH-sponsored portal for flow data at https://www.immport.org/home. Raw data and complete MS data sets for metabolomics have been uploaded to the Center for Computational Mass Spectrometry, MassIVE (link: ftp://MSV000098244@massive-ftp.ucsd.edu (MassIVE ID number: MSV000098244)). Cut&Run seq, DNA methylation and RNA-seq are available in GEO: link: https://www.ncbi.nlm.nih.gov/geo). Accession Number: Cut&Run seq (GSE310773), DNA methylation (GSE311289), and RNA-seq (GSE297039),

## Ethical Statement

All human blood donor samples were purchased from Vitalant (San Francisco, CA). All materials were fully de-identified prior to receipt by the investigators except information regarding age, sex and CMV serostatus. According to institutional guidelines and federal regulations (45 CFR 46) commercially supplied de-identified human blood products do not constitute human subjects research and is exempt from IRB review. No identifiable donor information was available to the research team except age, sex and CMV serostatus. All procedures complied with the ethical standards of the Buck Institute and the guidelines of the Declaration of Helsinki.

## Notes

### Competing Interest Statement

The authors have declared no competing interest.

### Summary of Updates

This version includes revised funding information.

